# Effect of cutting depth during sugarcane harvest on root characteristics and yield

**DOI:** 10.1101/2020.08.11.246033

**Authors:** Yue-bin Zhang, Shao-lin Yang, Jun Deng, Ru-dan Li, Xian Fan, Jing-mei Dao, Yi-ji Quan, Syed Asad Hussain Bukhari, Zhao-hai Zeng

## Abstract

Ratooning is an important cultivation practice in sugarcane production around the world, with underground buds on the remaining stalk acting as the source for establishment of a subsequent ratoon crop. However, the optimal depth of cutting during harvest in terms of yield and root growth remains unknown. We carried out a two-year field study to determine the effects of three cutting depths (0, 5 and 10 cm below the surface) ratoon cane root and yield. Results showed that cutting to a depth of 5 cm increased the root fresh weight and root volume by 32-40% and 49-85%, respectively, compared to cutting depths of 0 and 10 cm. Remarkably, cutting to a depth of 5 cm also had a significant effect on the development of fine roots, which is closely linked to cane yield. The effect was particularly noticeable in terms of two root traits, root volume and the surface area of roots with a diameter of 1.0-2.0mm, and root length and the number of root tips in roots with a diameter of 0-0.5mm. As a result, a cutting depth of 5 cm below the surface increased cane yield by 35 and 25% compared to depths of 0 and 10 cm below the surface, respectively. Overall, these findings suggest that a cutting depth of 5 cm is optimal in terms of sugarcane yield, largely due to the enhanced effect on root traits, especially the development of fine roots. These findings will help optimize sugarcane ratoon management and improve the ratoon cycle.

## 1. Introduction

Sugarcane is an economically important crop in tropical and subtropical regions, and is currently cultivated in 121 countries around the world [1]. A large, perennial, tropical C_4_ grass, sugarcane stores sucrose in its stem, and has long been recognized as one of the most efficient crops at converting solar energy into chemical energy, harvestable as sucrose and biomass [2]. After harvest, the remaining underground buds generate shoots, which go on to establish a subsequent ratoon crop [3]. The main advantages of ratooning are the cost savings related to planting labor, purchase of cane setts and soil tillage [4,5]. A reduction in costs of 45% has been reported with an overall increase in net benefits [6]. Ratooning also results in early sprouting and ripening compared with newly planted crops [7]. Accordingly, 6 to 20 ratoon crops are often harvested from a single planting in many countries [5,8], representing more than 50% of the total sugarcane production in tropical regions and more than 40% in sub-tropical regions [9]. For example, ratoon crops account for about 55% of the sugarcane produced in India [10], 80-85% in Hawaii, 80-90% in Brazil, and at least 50% in China [11,12]. Maintaining ratoon crop yield and the ratoon cultivation period is therefore an important consideration of sugarcane production.

The length of the sugarcane ratooning cycle is crucial to production and depends on the cultivar, local soil and environmental conditions, management practices, and the production level. In general, the cycle is approximately 4-5 years in Brazil, 5-7 years in Cuba, and 5-8 years in the Mauritius [13]. However, in China, the cycle is relatively short, with only two ratoon crops [14] due to the sharp decrease in yield. Yield decline is mainly caused by improper ratoon management [3], with shallowing of the underground buds and a subsequent decline in activity followed by eventual failure to germinate.

A few studies have examined the effects of stubble height on yield in ratoon crops such as forage sugarcane [15,16], rice [17,18], and alfalfa [19], as well as the effect of stubble height after mowing [20,21]. However, little is known about the effects of cutting depth or the length of remaining stalk on ratoon cane yield or the ratooning cycle. An increase in stubble height was found to increase ratoon crop yield in forage sugarcane [16], while a decrease in stubble height caused an increase in ratoon crop yield in alfalfa [19], rice [17] and forage grasses [20,21]. Moreover, increased yield was found to correspond to increasing residual height of the second crop in rice [18].

Stubble shaving is a popular management practice in sugarcane ratoon crops [4,15,22], whereby some of the aboveground residue and terminal buds are removed [23]. However, compared to harvest, stubble shaving is thought to be harmful to the remaining stalks and below ground buds [15]. Furthermore, the optimal depth of stubble shaving has yet to be determined. According to Li [7], the length of cane stalk left below ground is more than 20 cm under conventional cultivation of new plant cane in China. However, with increased shavings, the position of the ratoon stool is thought to become shallower compared with the original cane sett. Accordingly, the root system from the ratoon stool then generates in these shallower soil layers, with no deep nodes from which new roots can develop, having an effect on ratoon yield and shortening the ratoon cycle. It has therefore been suggested that deeper harvesting might be more suitable in terms of root growth and shoot node development underground. Moreover, while the removal of upper layers is practical, the most beneficial cutting depth in terms of the germination of underground buds with a robust root system is a key consideration. In addition, although the effect of stubble shaving on ratoon crops has been examined, most of these studies are concerned with the remaining stubble and aboveground residues, with little attention being paid to the underground parts. Understanding the optimal depth of cutting during harvest and the effect on yield is therefore essential to sugarcane production.

Since underground buds are important in establishing ratoon crops [24], proper ratoon management aimed at promoting underground germination is important. The activity of underground buds was found to decrease, while the number of buds increases with soil depth [7]. In addition, as mentioned above, the root system of sugarcane plants is largely influenced by the depth of these underground buds [7]. Roots originating in deeper soil are stronger and have a greater mass compared with those generated in shallower soil [25]. Cultivation practices that promote root development in deeper soil should therefore result in a stronger root system [22], and subsequent growth of healthy tillers with strong anchorage [8]. We therefore hypothesized that deeper cutting would increase ratoon crop yield by stimulating the germination of buds at lower stalk positions belowground, thereby promoting root development on these nodes, and increasing the absorption of water and nutrients from deeper rainfed soil. We therefore varied cutting depth to examine changes to bud germination, root growth, and root morphology, and the effects of root development on ratoon yield. The findings will help optimize ratoon crop management and improve the ratoon cycle in this region.

## 2. Materials and methods

### 2.1. Site description

The field experiment was conducted in Kaiyuan City (103°15’E, 23°42’ N, 1117 m a.s.l.), Yunnan Province, Southwest China. The soil is sandy loam with 50.0% sand, 34.0% silt, and 16.0% clay, and a pH of 7.5. The soil has a high organic matter content of 1.8%, with a total potassium content of 0.76%, total nitrogen content of 0.142%, and total phosphorus content of 0.12%. The content of alkaline hydrolysis nitrogen is 78.2 mg kg^−1^, available phosphorus is 26.6 mg kg^−1^, and available potassium is 49.0 mg kg^−1^. Organic matter was determined using the potassium dichromate method, soil nutrients were tested using flame atomic absorption spectrophotometry, and soil mechanical composition was determined using the sedimentation method. The field experiment was conducted under rainfed conditions, with a groundwater level of approximately 2 m below the ground surface.

The climate in this region is characterized by hot summers and long winters, with severe droughts in early spring. The highest precipitation and temperatures occur in June and July. Mean annual precipitation is 983 mm, and winter and summer precipitation are 20.3 and 174.6 mm per month, respectively. The mean air temperature is 20.0 °C, and the mean winter and summer temperatures are 14.4 and 24.1 °C, respectively. These annual figures are also of intrinsic value since they behave differently with respect to each other during the growing season from June to October (Fig. 1).

**Fig. 1.**
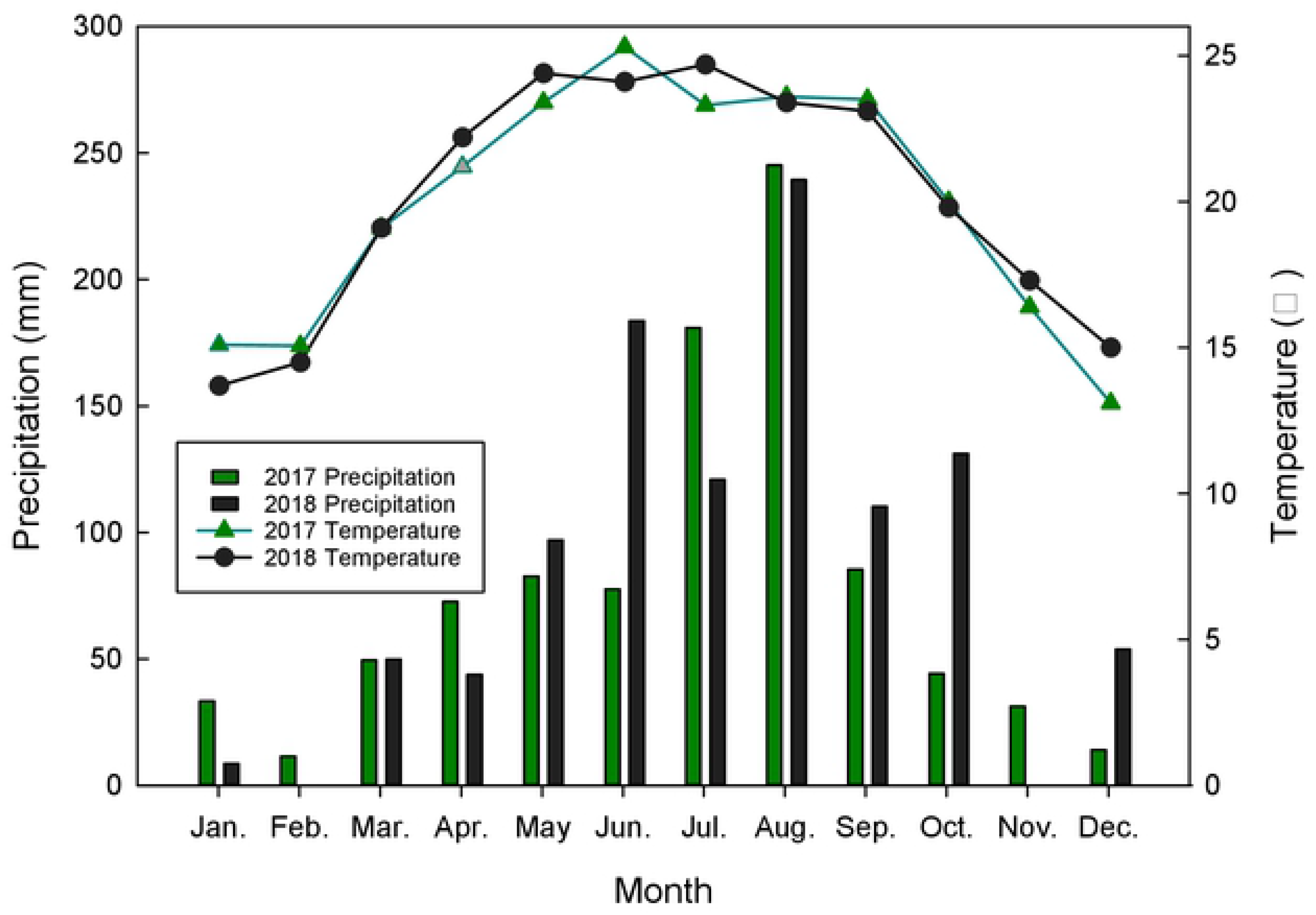
Monthly precipitation and temperatures at the experimental site (Kaiyuan City, Yunnan Province, southwest China) for 2017 – 2018.

### 2.2. Experimental design

The experiment was conducted from 2017 to 2018 using sugarcane cultivar YZ081609 [24]. Planting was carried out on 22 April 2016 at a conventional planting density of 120,000 buds ha^−1^ (two-bud cane setts). The planting depth (cane sett bed to soil surface) was approximately 10 cm, with about 10-15 cm of earthing up before the grand growth phase. Thus, at harvest, 20-25 cm of soil covered the original cane setts. NPK fertilizer (20: 12: 18) was applied during the early elongation period at a rate of 1200 kg ha ^−1^ on 10 March 2017 and 10 March 2018. All other cultivation and crop management procedures were in line with conventional cropping practices.

Based on the density of underground buds on the ratoon stool and the germination potential, we divided the underground buds into three types (Fig. 2). Type 1: terminal buds in the top soil. Least abundant, but fastest to germinate, with the roots distributed mainly in the upper soil layer. Type 2: distributed at a depth of 5-10 cm, mostly active and fast to germinate, with deeper roots than type 1. Type 3: distributed below 10 cm, mostly dormant with the poorest rate of germination. Accordingly, cutting treatment was carried out at the following depths: the soil surface (0 cm, T1), and 5 (T2) and 10 cm below the surface (T3) (Fig. 2). All plants were harvested manually using a sharp hoe. In order to maintain an accurate cutting depth, one side of the soil surrounding each stool was removed to the respective depth under each treatment before harvest. Plant crops (2016) were harvested on 9 March 2017, cultivated to the first ratoon crop then re-harvested on 4 March 2018, representing the second ratoon crop. Each plot was 30 m^2^, with five rows, each 6 m long. Each treatment was randomized and consisted of 3 replications.

**Fig. 2.**
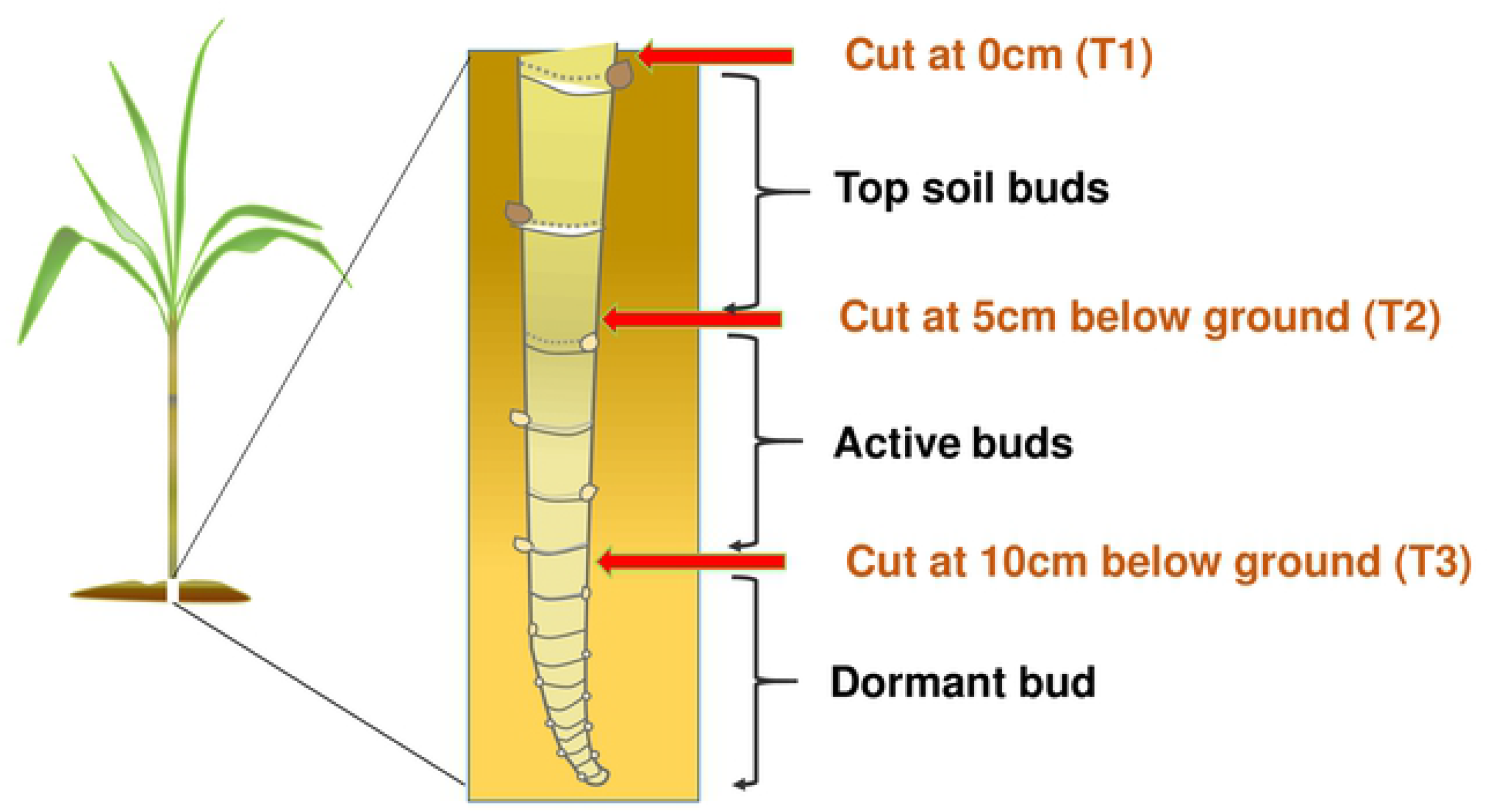
Schematic diagram of the experimental design.

### 2.3. Data collection

Root and shoot biomass were investigated at three stages in the first and second ratoon crops: the early stem stage of elongation (168 days after harvest, DAH), the middle stage of elongation (198 DAH), and the late stage of elongation (238 DAH). Three randomly selected stools from each plot were cut at ground level then their fresh weights were measured immediately. The shoots were then dried at 70 °C for 72 hours to measure the dry weight. The root system was excavated using a frame (50×50 cm) to a depth of 30 cm and the roots were washed thoroughly. After measuring the fresh weight, the roots were analyzed for the following morphological characteristics: length, surface area, volume, tip number, fork number, crossing (of growth in a three-dimensional space), and diameter using a root scanner (Win RHIZO 2009 b; REGENT Instruments Inc., Quebec, Canada). After scanning analysis, the roots were dried at 70 °C for 72 hours to measure the dry weight. The length, surface area, volume and tip number were also analyzed in terms of a range of root diameters to evaluate the impact of treatment on fine root development. At maturity (9 March 2017, and 4 March 2018), plant height and cane diameter were measured in 10 random selected plants in each plot then the number of millable stalks and the overall yield were determined for each plot.

The economic benefits of each cutting depth were also calculated based on the resulting inputs (¥): 120 Yuan ton^−1^ for harvest, 2,250 Yuan ha^−1^ for trash pulverizing, 4,800 Yuan ha^−1^ for fertilizer, 750 Yuan ha^−1^ for fertilizer application and hilling up, 900 Yuan ha^−1^ for pesticide application, and 450 Yuan ton^−1^ for output of millable cane, in 2017 and 2018, respectively. Costs were calculated based on the area of the plots and the cultivar used.

### 1.4. Statistical analyses

When the P value of the correlation between root biomass and dry weight or shoot biomass and dry weight was greater than that of the correlation with fresh weight, the dry weight was used for analysis. In all other cases, the fresh weight was used. Mean differences in sugarcane yield and yield components were compared separately each year using one-way analysis of variance (ANOVA) according to the Tukey method with treatment as the fixed effect and replication as the random effect. The shoot and root biomass, and root traits of each plant were averaged for each cane cluster (amount per cluster / millable cane per cluster). Mean differences in shoot and root biomass, root volume, root surface area, root length, and root tip number were then compared separately for each year using one-way ANOVA according to the Tukey method, with treatment as the fixed effect and replication as the random effect (SPSS 19.0 statistical software and Microsoft Office Excel). Correlation analyses were carried out using the Bivariate Correlation Method with Pearson’s correlation coefficient (SPSS 19.0 statistical software). The relationships between root traits and shoot biomass were determined using Microsoft Office Excel 2016. We also estimated the root traits (root fresh weight, root volume, root surface area, root length, root tip number, root fork number, root crossings) per square meter and at a soil depth of 0.3 m according to the number of millable canes per unit area.

## 3. Results

### 3.1. Yield and benefits

Cutting depth had a significant effect on cane yield in both the first and second ratoon crops, with T2 (cutting at 5 cm below the surface) performing significantly better than the other two cutting depths. Stalk diameter was also greatest under T2, while plant height decreased in both ratoon crops with increasing cutting depth during harvest. The number of millable stalks increased with increasing cutting depth, with an increase from T1 to T2 in both ratoon crops, but no significant increase from T2 to T3 (Table 1). Cutting depth also had a significant impact on the millable cane number, stalk height and stalk diameter. Overall, stalk diameter is thought to be the main factor affecting the difference in yield, with stalk height and the number of millable canes acting as composite factors.

**Table 1.**
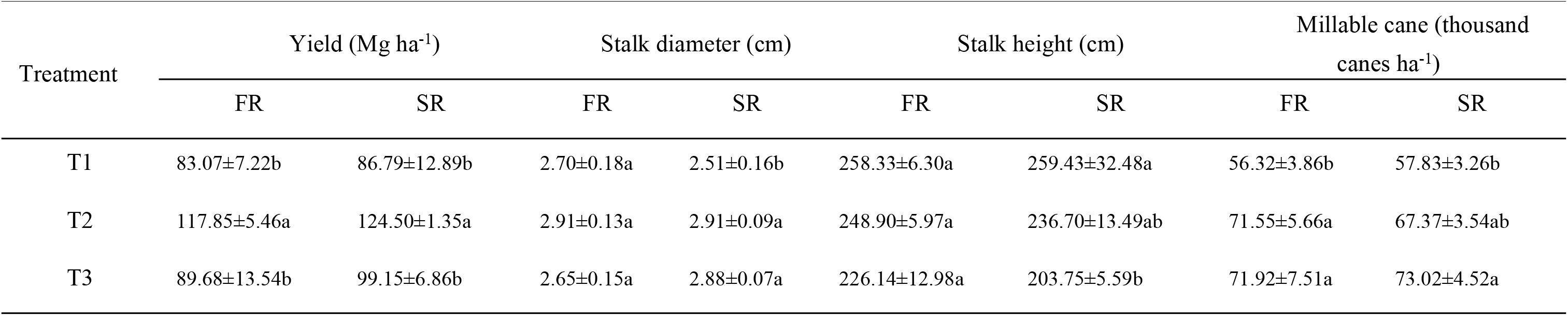
Sugarcane yield and yield components at cutting depths of 0 cm (T1), and 5 (T2) and 10 cm below the ground (T3) in the first (FR) and second ratoon crop (SR). Different letters in the same column indicate statistical significance (one-way ANOVA Tukey, P < 0.05).

The net economic benefits were significantly impacted by cutting depth in both ratoon crops, at 16,800, 27,700, and 19,600 Yuan ha^−1^ under T1, T2 and T3, respectively. Moreover, T2 resulted in an increase in net benefits of 10,900 Yuan ha^−1^ (64.8%) and 8,000 Yuan ha^−1^ (40.9%) compared to T1 and T3, respectively, in both ratoon crops (Fig. 3.).

**Fig. 3.**
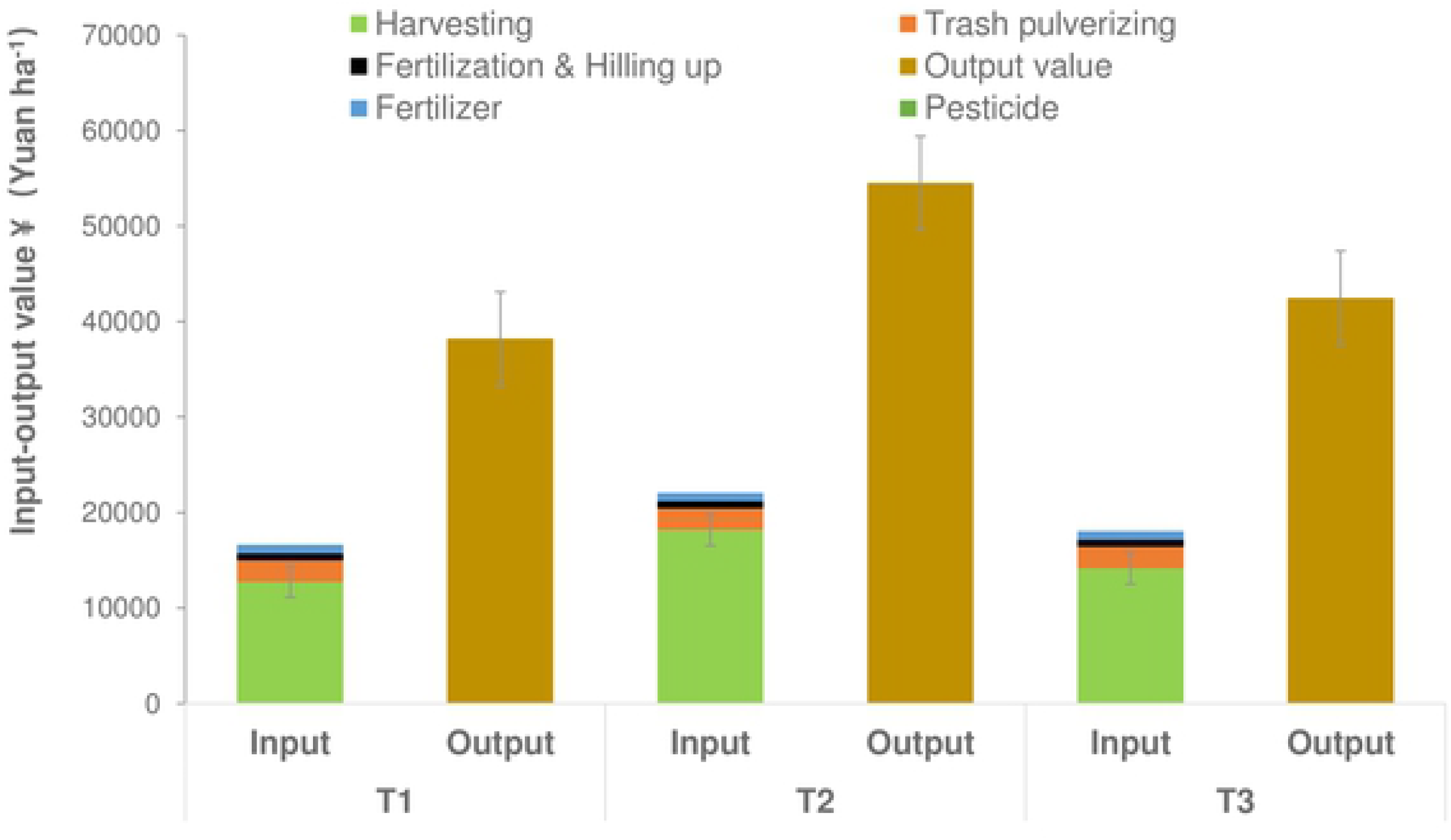
Mean input-output values of the two ratoon crops at each cutting depth (0 cm (T1), 5 cm below the ground (T2), and 10 cm below ground (T3)). The net benefits of each treatment were calculated based on the following inputs in RMB(¥): 120 Yuan ha^−1^ for harvesting, 2,250 Yuan ha^−1^ for trash pulverizing, 4,800 Yuan ha^−1^ for fertilizer, 750 Yuan ha^−1^ for fertilizer application and hilling up, 900 Yuan ha^−1^ for pesticides, and 450 Yuan ton^−1^ for output of millable canes. There was little change in the input price between study years. Costs were calculated according to the area size and the sugarcane cultivar. Error bars, SD.

### 3.2. Relationship between root and shoot traits

#### 3.2.1. Root fresh weight and shoot biomass under each cutting depth

Cutting depth had a significant impact on shoot and root biomass in both ratoon crops, especially the second (Fig. 4). Shoot biomass and root fresh weights followed a consistent trend, increasing then decreasing in both the first and second ratoon crops, and were highest under T2 at all three elongation stages (Fig. 4a, 4b). The highest root fresh weight was also observed under T2 compared to T1 and T3, and a similar trend was observed in terms of the shoot fresh biomass and root fresh biomass on the same DAH, especially in the second ratoon crop. Overall, there was little difference between the changes in shoot and root biomass between the two ratoon crops, with an increase in shoot biomass and a decrease in root biomass with increasing DAH in the first ratoon crop.

**Fig. 4.**
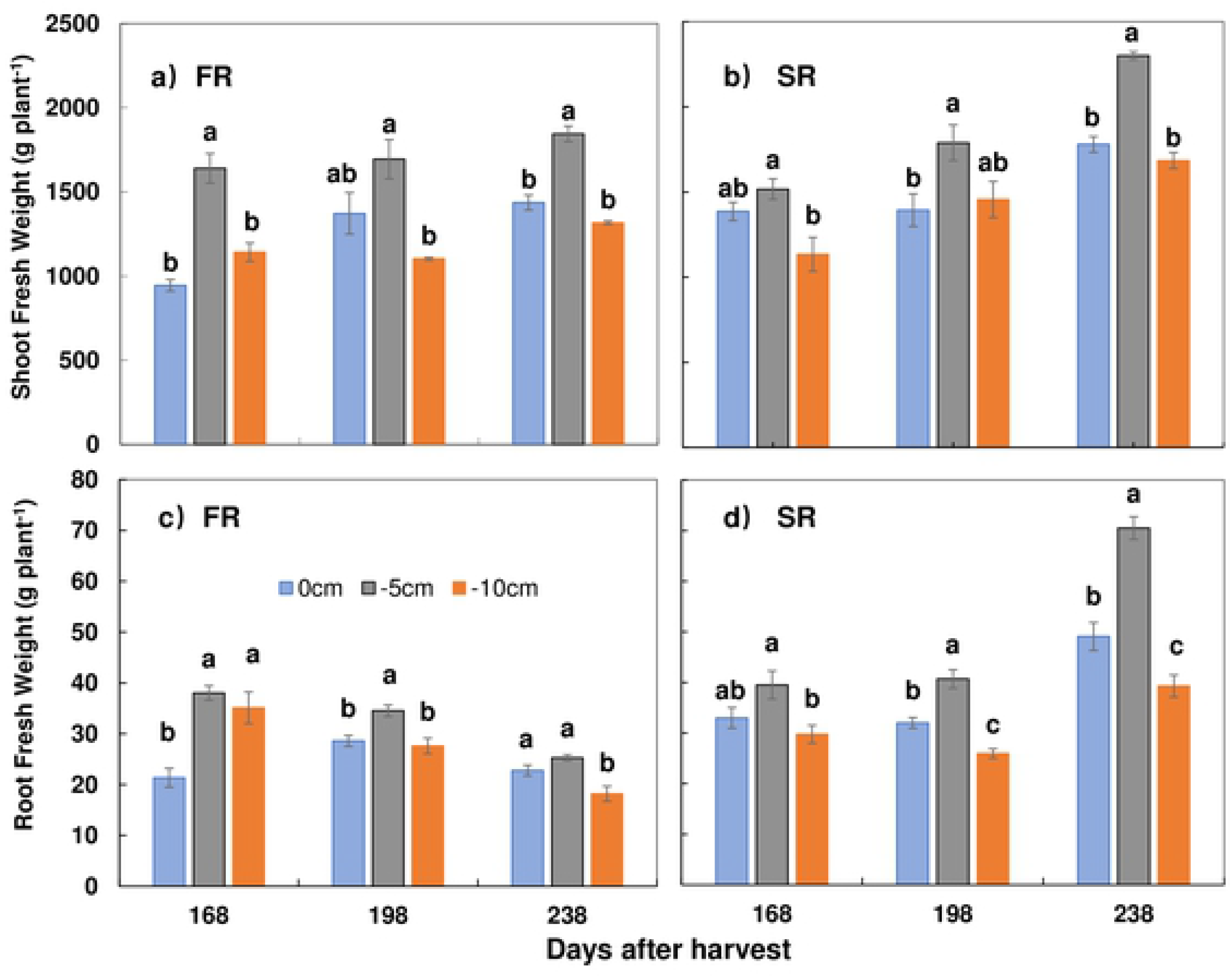
Shoot and root fresh weight under each cutting depth in the first (FR) and second ratoon crops (SR). Days after harvest (DAH) of 168 to 238 represent three months of elongation, while 238 DAH represent the end of the elongation stage, approaching maturity stage, with a slow increase in yield. g plant^−1^: the amount per each millable cane in a cluster. Different lower-case letters indicate significant differences within treatments at a significance level of P < 0.05 level in each year. Error bars, SD.

### 3.2. Relationship between root morphological characteristics and shoot biomass

Due to the similar trends in shoot and root biomass at each stage in both ratoon crops (Fig. 3), correlation analysis was carried out between shoot fresh weight and root morphology. As a result, shoot biomass was found to be positively correlated with root biomass, root length, root surface area, root volume, root tip number, root fork number and root crossings in both ratoon crops (P < 0.01). Moreover, the r values of these correlations in the first and second ratoon crops respectively were in the order of root biomass (0.86 and 0.78) > root fork number (0.77 and 0.65) > root volume (0.67 and 0.72) > root length (0.68 and 0.72) > root crossings (0.77 and 0.59) > root tip number (0.62 and 0.69) > root surface area (0.70 and 0.56) (Table S1).

A linear relationship was observed between the root morphological characteristics and shoot biological yield in both ratoon crops, and between the root fresh weight and shoot fresh biomass, the root volume and shoot fresh biomass, and the root fork number and shoot fresh biomass (Fig. 5). R^2^ values were much lower in the first ratoon crop than the second crop, and the scatterplots of the first crop were much more concentrated than in the second crop, but the correlation coefficient (r) was extremely significant (P<0.01) as shown in Table S1. Moreover, the root tip number and fresh shoot biomass, the root length and fresh shoot biomass, and the root surface area and fresh shoot biomass all showed the same patterns (Fig. S1). This was not because the fresh weight of the first year’s shoots was poorly correlated with the root parameters; on the contrary, the correlations were extremely significant in the first ratoon, but because the distribution of the shoot fresh weight was more concentrated with the root parameters compared with the second ratoon crop. This may be attributed to the cumulative effect of the stalk cutting position from the first ratoon crop to the second, and also the difference in rainfall in June and July between the two seasons.

**Fig. 5.**
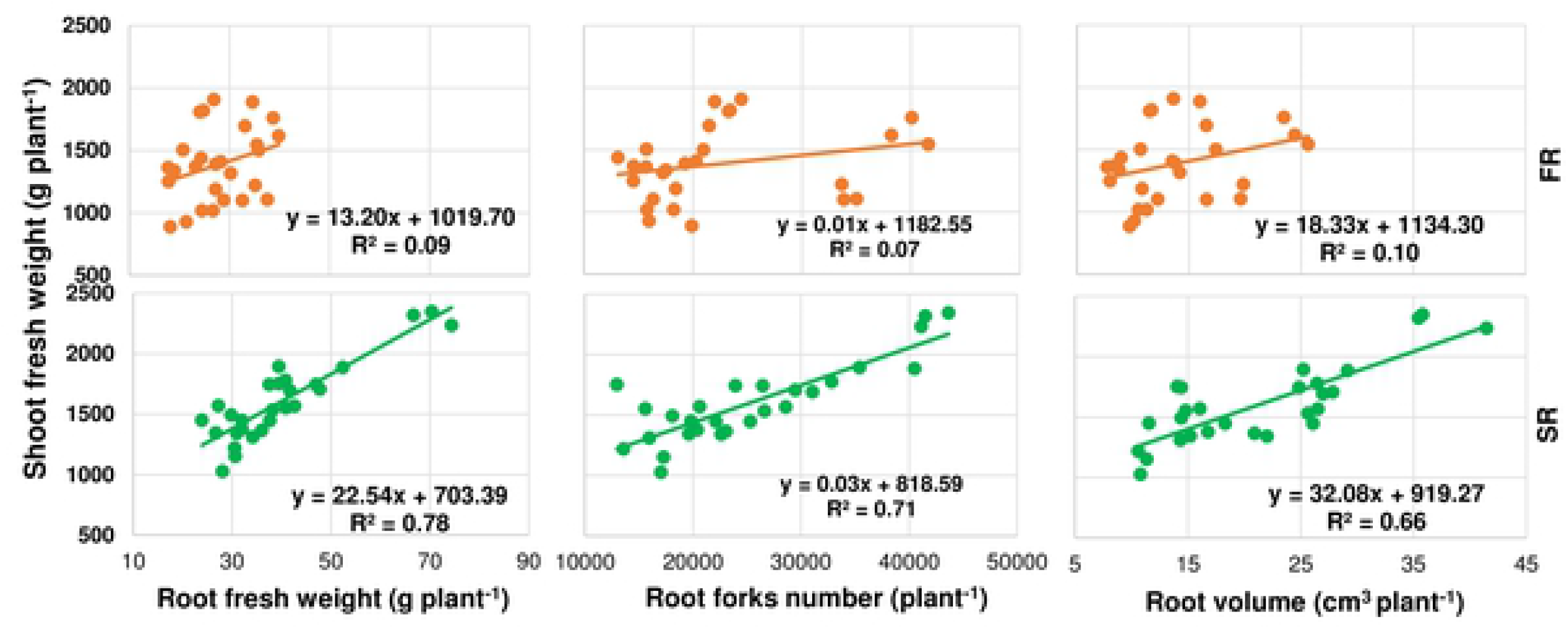
Relationships between shoot biomass and root fresh weight, root fork number, and root volume in the first (FR) and second ratoon crop (SR). Plant^−1^: the amount per each cluster / millable canes in the cluster.

### 3.3. Root characteristics under each cutting depth according to root diameter

The results showed that the greatest root volume and root surface area were observed in roots with a diameter of 1.5-2.0 and 1.0-1.5 mm, respectively. Moreover, values were greater under T2 than T1 and T3. The greatest root length and root tip number were observed in roots with a diameter of 0-0.5 mm, with root length decreasing much slower than the root tip number as the root diameter increased. Root volume and root surface area were therefore considered a similar morphological index, since values reached a maximum at a middle root diameter of 1.0-2.0 mm. Meanwhile, root length and root tip number reached a maximum in roots with the finest diameter (Fig. 5,6).

**Fig. 6.**
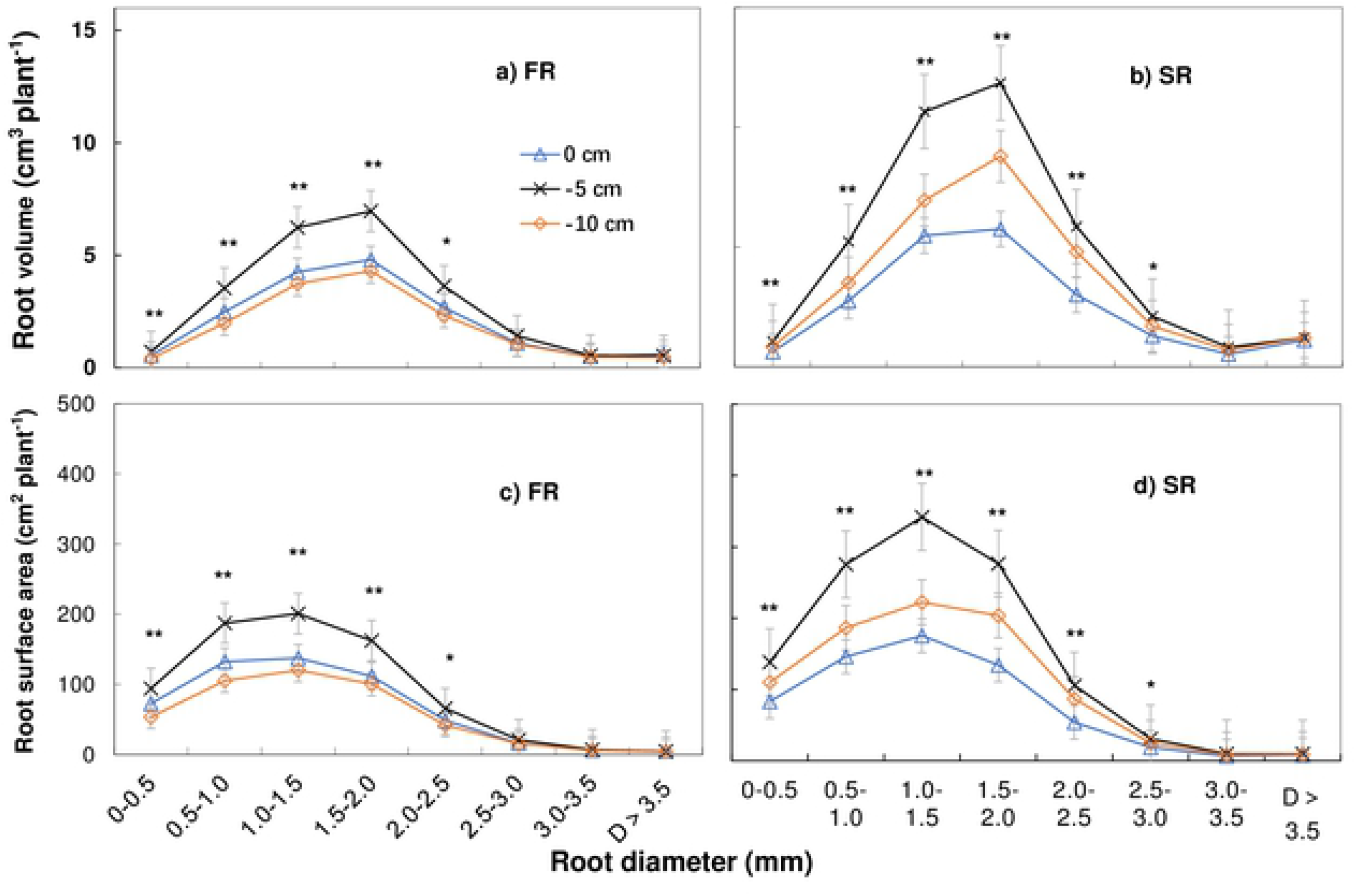
Effect of cutting depth on root volume and root surface area according to root diameter (D) in the first (FR) and second ratoon crop (SR). Root volume was highest in roots with a diameter of 1.5-2.0 mm in both FR and SR, while the root surface area was highest at a diameter of 1.0-1.5 mm plant^−1^: the amount in each cluster/ millable canes in the cluster. * P < 0.05, ** P < 0.01. Error bars, SD.

The average root diameter was similar under all three cutting depths, at 0.87, 0.89 and 0.90 mm under T1, T2 and T3, respectively. Each root trait showed differences among treatments within different root diameter ranges, although there were only slight differences among treatments in roots with a diameter of 0-3.5 mm (Table 2). That is, the treatments did not change the percentage of each part of root in each plant. In terms of root volume, 25.9%, 29.4%, and 15.6% of roots had a diameter of 1.0-1.5, 1.5-2.0 and 2.0-2.5 mm, respectively. The root surface area of roots with a diameter of 0.5-1.0, 1.0-1.5 and 1.5-2.0 mm was 23.7, 27.1 and 22.3%, respectively. Similarly, the root tip number in roots with a diameter of 0-0.5 mm was 90.2%, while the root length was 40.9%. The quantity of each trait in a given root diameter was therefore influenced by cutting depth and growing year, although the overall quantitative ratio differed slightly among treatments.

**Table 2.**
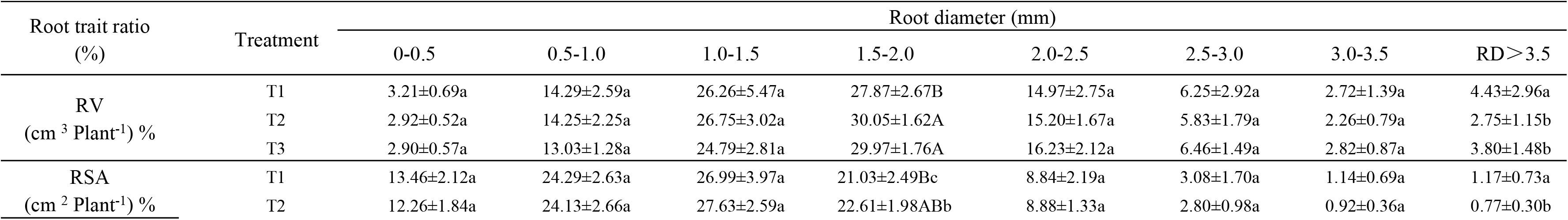
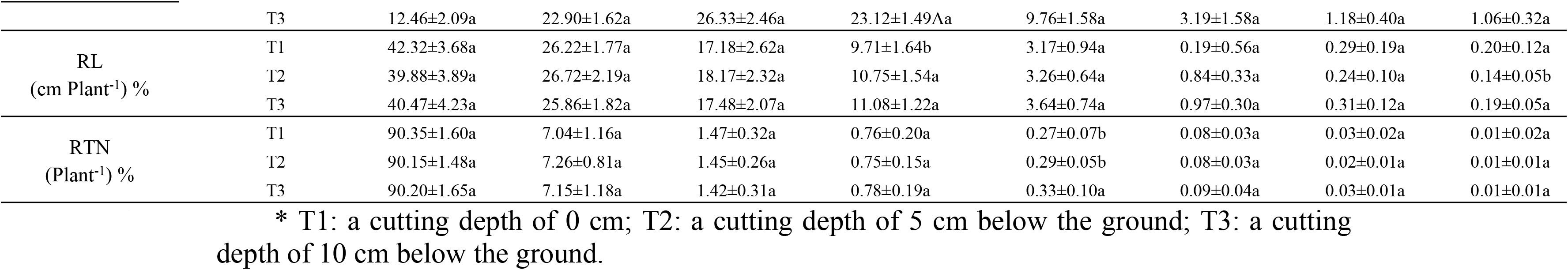
Root volume (RV), root surface area (RSA), root length (RL) and root tip number (RTN) according to root diameter (RD). ± represents the standard deviation. Different lowercase letters in the same column under each cutting depth denote a significant difference at P < 0.05, while different uppercase letters denote a significant difference at P < 0.01.

We also estimated the root traits per square meter and a 0.3 m soil depth. Accordingly, the root fresh weight ranged from 167-246 g m^−2^, root volume from 79-164 m^3^ m^−2^, root surface area from 3628-7329 cm^2^ m^−2^, root length from 13,430-18,390 cm m^−2^, root tip number from 36,538-68,473 m^−2^, root forks number from 114,364-215,035 m^−2^, and root crossings from 9,059-12,038 m^−2^. The amount of root forks was highest, followed by the number of root tips, possibly because of the specific morphology of the root tip, the diameter of which decreases in a fitted ellipse [26]. Some root tips will also have been destroyed or damaged during harvest.

### 3.4. Root morphological traits are affected most by cutting depth

#### 3.4.1. Effect of cutting depth on root volume and root surface area according to root diameter

The analysis of root morphology revealed that root volume was highest in roots with a diameter of 1.5-2.0 mm in both ratoon crops, with significantly higher values under T2 (Fig. 6 a, b). Root volume under T2 reached 6.98 and 11.85 cm^3^ in the first and second ratoon crop, respectively, in roots of 1.5-2.0 mm, 2.17 and 3.06 cm^3^ higher than under T1 and T3, respectively (P < 0.01). A significantly higher root volume was also observed under T2 in roots with a diameter of 0-2.5 mm. However, in the first ratoon crop, no significant differences were observed among treatments in roots with a diameter > 2.5 mm, and at a diameter of > 3.0 mm in the second ratoon crop.

The root surface area was highest in roots with a diameter of 1.0-1.5 mm in both ratoon crops, with significantly higher values under T2 (P < 0.01). Moreover, a significantly higher root surface area was also observed under T2 in roots with a diameter of 0-2.5 mm. Meanwhile, no significant differences were observed between treatments in roots > 2.5 mm in the first ratoon crop and > 3.0 mm in the second ratoon crop (Fig. 6 c, d). Under T2, the root surface area reached 200.8 and 341.75 cm^2^ plant^−1^ in roots with a diameter of 1.0-1.5 mm in the first and second ratoon crops, respectively, 63.6 and 119.42 cm^2^ plant^−1^ higher than under T1 and T3, respectively.

A higher root volume and root surface area were therefore observed under T2 compared to T1 and T3 in roots of different diameters, especially in the second ratoon crop, with an increase in the maximum value as well as the total amount. A significant change was also observed at a root diameter of 2.0-2.5 and 2.5-3.0 mm. Overall, between the first and second crops, only slight increase was observed in T1 compared to T1 and T3 in terms of both root volume and root surface area.

#### 3.4.2. Effect of cutting depth on root length and root tip number according to root diameter

Cutting depth also affected the root length and root tip number in both ratoon crops. A significant increase in both parameters was observed under T2 in roots with a diameter < 2.5 mm, while no differences were observed at a diameter of 3.0-4.5 mm. Values were highest in roots with a diameter of 0-0.5 mm (Fig. 7) and were significantly higher under T2 than the other treatments in both ratoon crops (P < 0.01). Overall, both the root length and root tip number were greater in T2 than T1 and T3 at different root diameters, especially in the second ratoon crop, with increase in the maximum value as well as the total amount. Moreover, a significant change was also observed at a diameter of 2.0-2.5 mm and 2.5-3.0 mm. Between the first and second crop, only a slight increase was observed in T1 compared to T3 in terms of both root length and root tip number.

**Fig. 7.**
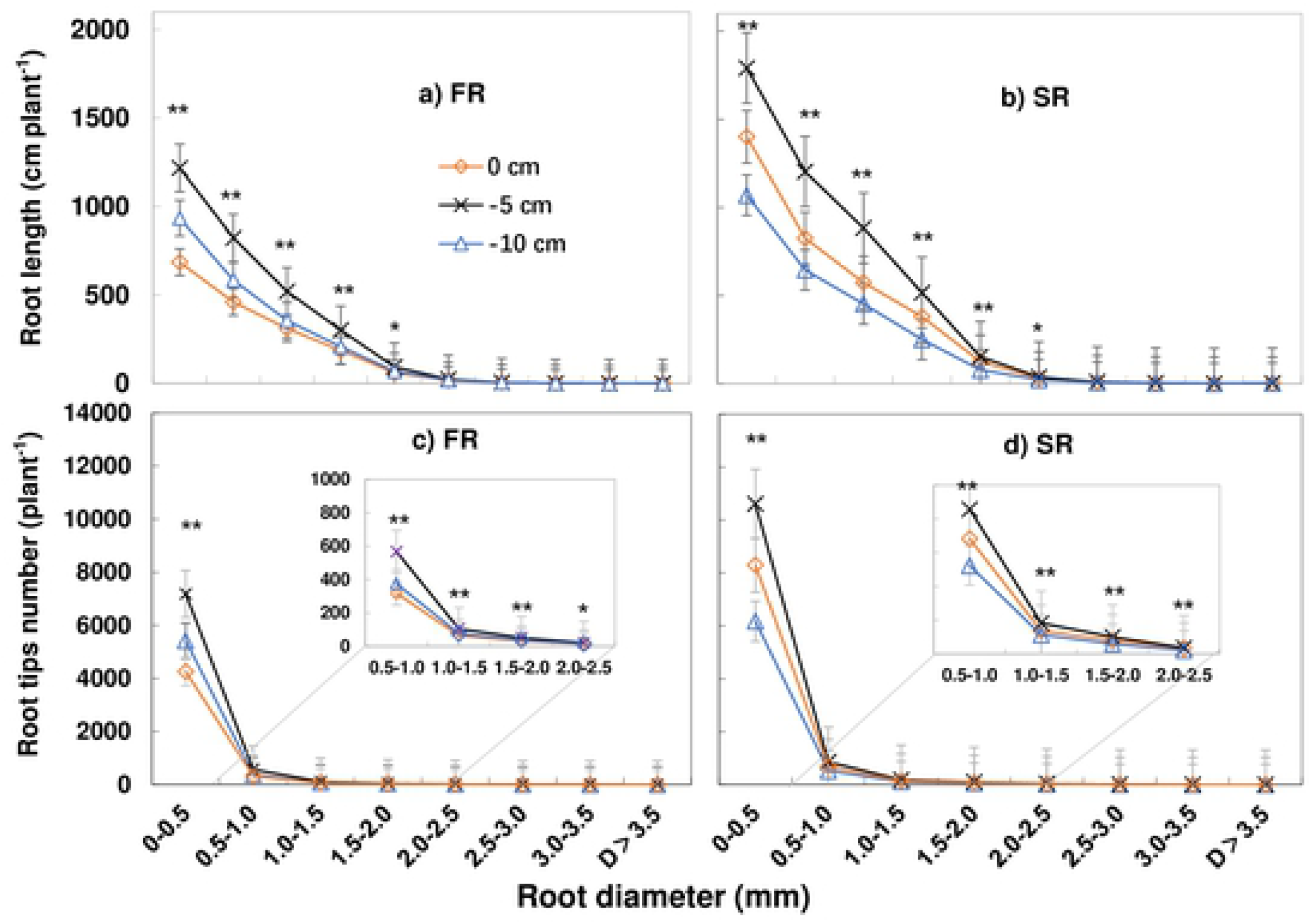
Effect of cutting depth on root length and root tip numbers. At the diameter of 0~0.5mm, the root length and root tips number of each plant reached the highest, in both FR and SR. /plant: the amount of each cluster/ millable canes of the cluster. * P < 0.05. ** P < 0.01. Error bars, SD.

The longest root length and maximum number of root tips was also determined in roots with a diameter of 0-0. 5 mm (Fig. 7.), with a clear decrease in root tip number compared to root length at diameters > 0.5 mm. Under T2, the root length was 1218 and 1790 cm plant^−1^ in roots with a diameter of 0-0.5 mm in the first and second ratoon crop, 283 and 389 cm plant^−1^ greater than under T1 and T3, respectively. Moreover, the root tip number was 7184 and 10,608 plant^−1^ at a diameter of 0-0.5 mm in the first and second crops, 1768 and 2315 plant^−1^ more than under T1 and T3, respectively.

#### 3.4.3. Relationship between fine root characteristics and shoot biomass

The above results show that cutting depth affected the root biomass, and the values of four root components (root volume, root surface area, root length and root tip number) in roots within a diameter range of 0-2.5 mm. However, there was little effect on any of the four root components in roots with a diameter > 2.5. We therefore analyzed the relationship between shoot biomass and the above root components.

The results showed better linear relationships than under the total range of root diameters between the root morphological characteristics and shoot biomass in both ratoon crops. Clear linear relationships were observed between the root volume and shoot fresh biomass, root surface area and shoot fresh biomass, root length and shoot fresh biomass, and root tip number and shoot fresh biomass in roots with a diameter of 0-2.5 mm (Fig. 8). The R^2^ values were much lower in the first ratoon crop compared to the second, and the scatterplots were more concentrated. Although the above relationships all showed a similar pattern, the R^2^ values were greater between the fine root volume and root surface area plus the shoot fresh weight. This may be attributed to the cumulative effect of stalk cutting position from the first ratoon crop season to the second.

**Fig. 8.**
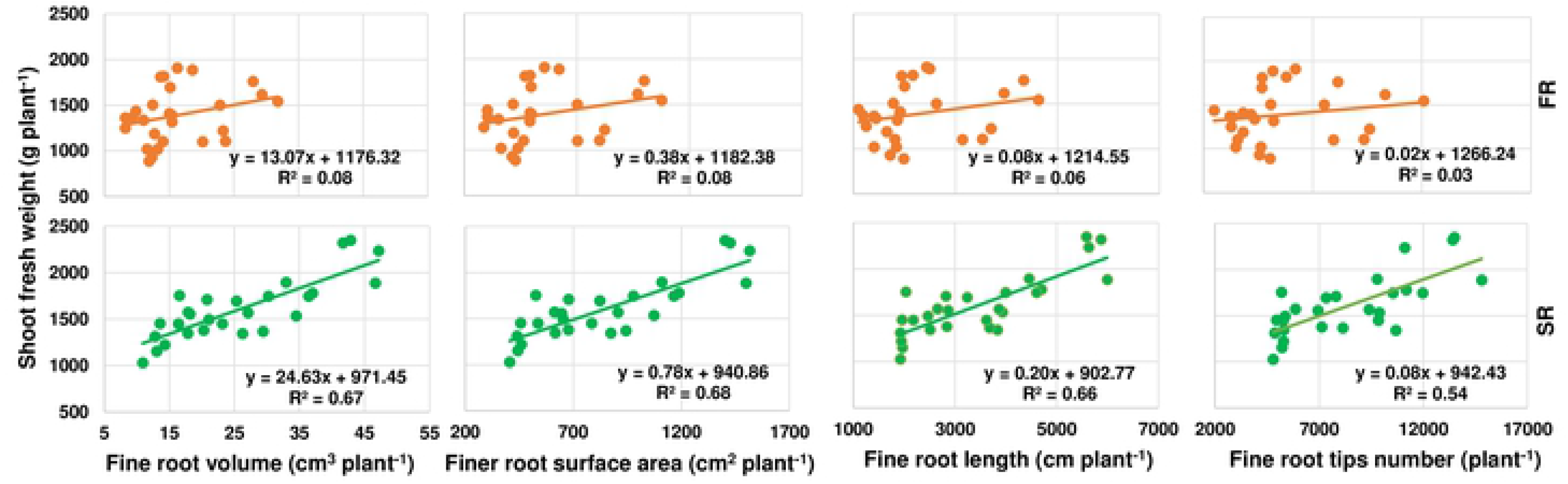
Relationships between shoot biomass and fine root characteristics in the first (FR) and second ratoon crop (SR). Plant^−1^: the amount per each cluster / millable canes in the cluster.

## 4. Discussion

Plant roots are strongly linked with morphological, physiological and biochemical functions [27]. Sugarcane roots perform a vital role in regulating both shoot development and responses to the environment [28]. Root morphology determines the ability of crop plants to explore the rhizosphere and efficiently absorb water and nutrients from the soil [29,30]. Root length, root surface area, and root volume are therefore commonly used to evaluate the performance of a root system [31,32]. In this study, root development was significantly impacted by cutting depth. Cutting to depth of 5 cm below the surface (T2) increased activity of buds and deeper roots, enhancing the ability to absorb nutrients and water from the soil, and thereby increasing germination and root development. In contrast, cutting at a depth of 0 cm (T1) resulted in shallow root development and a poor ability to absorb nutrients and moisture, while cutting to 10 cm below the surface (T3) resulted in mostly dormant buds, whose poor activity resulted in relatively poor root development.

Morphological attributes of the fine roots affect nutrient and water uptake as well as biomass accumulation [33], and the length and diameter of fine roots play a pivotal role in nutrient and water acquisition [34–36]. In this study, roots with a diameter of 0-2 mm comprised about 54% of the total root length, about 69% of the root volume, and 73% of the root surface area. Meanwhile, roots with a diameter of 0-0.5 mm represented about 90% of the root tip number, while those with a diameter > 2mm contributed to only a small proportion. The proportion of fine roots in sugarcane differs from that in wheat and pulse crops. For example, in pulse crops, roots with a diameter of 0-0.2mm comprise about 60% of the total root length [37]. The classification of fine roots also differs between crops and, in general is characterized by roots with a diameter of 0-0.2 mm in rice [38,39], wheat, and pulse crops [37]. However, in some species, such as poplar, switch grass, cool-season pasture grasses, corn, soybean [40], cotton, maize and sorghum [41], the difference between fine and thick or coarse roots is classified by a diameter of 2 mm. In terms of the proportion of roots (Table 2) and amount of roots (Figs. 5 and 6), the root volume, root surface area and root length of roots with a diameter of 0-2.5mm played an important role in this study, while root tip number was important in roots with a diameter of 0-0.5 mm.

Positive correlations were also observed among root traits and shoot biomass, with traits in roots with a diameter of 0-2.5 mm having the greatest impact on shoot biomass. We therefore removed the data of root traits in roots with a diameter > 2.5 mm, and reanalyzed the linear relationships between shoot biomass and root volume, root surface area, root length, and root tip number in those with a diameter of 0-2.5 mm. As a result, we found that R^2^ values were highest at a diameter of 1.5-2.0 mm in both ratoon crops. Moreover, there were significant differences in root traits at a diameter < 2.5 mm, and the number of root tips was significantly different at a root diameter < 0.5 mm. As the root diameter increased, the differences among cutting depths rapidly declined, suggesting that the fine roots are essential for growth and development of the underground root system in perennial sugarcane. That is, the more the number of fine roots, the greater the absorption of water and nutrients, and the higher the yield.

Studies suggest that sugarcane stools contain about 10-19 underground buds, and together with the roots, these underground buds and stalks combine to form the ratoon stool or crown [42–46], distributed about 15 cm below the soil surface. The upper section of the sugarcane stubble is mainly composed of surface buds, with about 2-3 living buds. After germination, the roots remain in shallow soil, and nutrient and water absorption is poor. The mid-section is the active bud section, with about 4-5 active buds. After germination, the roots extend deep into the soil, improving nutrient and water absorption, and promoting survival under drought conditions. The lower buds are mainly dormant, and tend to remain so. Relatively deep cutting is therefore thought to eliminate apical dominance of the upper buds and promote the germination of dormant buds on lower underground stalks.

In this study, the position of the terminal buds decreased with increasing cutting depth from 0 to 10 cm. As a result, germination of these lower buds resulted in more nodes for root development, which, in turn, increased opportunities for water and nutrient absorption compared to germination of the upper buds. Thus, the shoot and root biomass were greater under T2 than T1. Theoretically, the shoot and root biomass should also have been greater under T3 compared to T2; however, the opposite was observed. One reason for this is thought to be the time required to come out of dormancy and for development of active buds, with the majority of buds at this depth remaining dormant as in *Betula pendula* [47]. This is supported by a previous study by Harrell et al. [17] in ratoon rice, whereby a decrease in stubble height from 40 to 20 cm affected growth of the ratoon crop by shifting the panicle point of origin during early growth and delaying maturity. In our study, plant height was lowest under T3 (Table 1); however, the amount of millable cane was highest, and the diameter was greater than under T1. This also suggests that lower terminal buds benefit from a better growth environment after germination, although the drawback is delayed growth of aboveground shoots. It was previously reported that sugarcane stems originating from deeper soil produce a stronger root system [25], which is the basis of a good crop stand. In this study, T2 resulted in the highest root biomass, which in turn, allowed sufficient uptake of nutrients and water, increasing the shoot biomass compared with other treatments.

Overall, the findings of this study suggest that a cutting depth of 5 cm below the soil surface promotes growth the root system, especially in terms of the root volume, root surface area and root length of roots with a diameter of 0-2.5mm, thereby significantly improving the shoot biomass and cane yield. This cultivation practice is a simple addition to ratoon cutting management, increasing overall production of ratoon cane. This could therefore be applied successfully to rain fed slopes in China and other Southeast Asian countries in line with manual harvesting techniques. Meanwhile, in terms of mechanical harvesting, improvements could also be made to the cutting machines, such as installing a plow tooth above the disc blade to allow removal soil with the cane stalk or lowering of the disc blade to reduce abrasion. In line with this, technology has already been extended to a production area of 25,000 hectares in the sugarcane district of Dehong State, Yunnan Province, China, in 2018.

## 5. Conclusions

Deep cutting at depths of 5 and 10 cm below the soil surface (T2 and T3) considerably increased yield compared to traditional cutting at the soil surface (0 cm, T1). The highest yield increase was observed under T2, with an increase in production of 32.74 (34.9%) and 25.5 ton ha^−1^ (25.2%) compared to T1 and T3, respectively. These results suggest that the improvements in all root traits across all root diameters supported a greater increase in shoot biomass and yield. This was especially true in terms of root volume and root surface area in roots with a diameter of 0-2.5 mm, and root length and root tip number in those with a diameter of 0-0.5 mm, both of which also showed a good linear relationship. Moreover, since buds at the soil surface and dormant buds remained under T1 and T3, this may have led to a shallower stool and delayed vegetative growth of the shoot, respectively. Cutting at a depth of 5 cm below the soil surface is therefore recommended in terms of root biomass, subsequent shoot biomass, and overall sugarcane yield.

## Acknowledgments

The study was funded by National sugar industry technical system (CARS-170205).

---

**Table S1.** Correlation between root morphology and shoots biomass. RFW: root fresh weight; RDW: root dry weight; SFW: Shoot fresh weight; SDW: Shoot dry weight; RL: root length; RSA: root surface area; RAD: root average diameter; RLPV: root length per volume; RV: root volume. First ratoon crop (FR) n=53, second ratoon crop (SR) n=43. ** P<0.01, * P<0.05 (2-tailed).

|  | Ratoon | RFW(g) | RDW(g) | SFW(g) | SDW(g) | RL<br>(cm) | RSA<br>(cm<sup>2</sup>) | RAD<br>(mm) | RLPV<br>(cm) | RV<br>(cm<sup>3</sup>) | Tips | Forks | Crossings |
| --- | --- | --- | --- | --- | --- | --- | --- | --- | --- | --- | --- | --- | --- |
| RFW(g) | FR | 1 | 0.967** | 0.859** | 0.841** | 0.836** | 0.875** | -0.107 | 0.155 | 0.893** | 0.812** | 0.830** | 0.792** |
|  | SR | 1 | 0.945** | 0.782** | 0.782** | 0.841** | 0.754** | 0.098 | -0.323* | 0.846** | 0.811** | 0.756** | 0.689** |
|  | - | - | -0.022 | -0.077 | -0.059 | 0.005 | -0.121 | 0.205 | -0.478 | -0.047 | -0.001 | -0.074 | -0.103 |
| RDW(g) | FR | 0.967** | 1 | 0.804** | 0.791** | 0.783** | 0.845** | 0.003 | 0.065 | 0.889** | 0.776** | 0.761** | 0.718** |
|  | SR | 0.945** | 1 | 0.779** | 0.779** | 0.799** | 0.706** | 0.137 | -0.348* | 0.809** | 0.764** | 0.737** | 0.684** |
|  | - | -0.022 | - | -0.025 | -0.012 | 0.016 | -0.139 | 0.134 | -0.413 | -0.08 | -0.012 | -0.024 | -0.034 |
| SFW(g) | FR | 0.859** | 0.804** | 1 | 0.979** | 0.683** | 0.702** | -0.142 | 0.180 | 0.703** | 0.618** | 0.767** | 0.766** |
|  | SR | 0.782** | 0.779** | 1 | 10.000** | 0.722** | 0.559** | 0.110 | -0.183 | 0.716** | 0.686** | 0.654** | 0.591** |
|  | - | -0.077 | -0.025 | - | 9.021 | 0.039 | -0.143 | 0.252 | -0.363 | 0.013 | 0.068 | -0.113 | -0.175 |
| SDW(g) | FR | 0.841** | 0.791** | 0.979** | 1 | 0.657** | 0.672** | -0.165 | 0.185 | 0.670** | 0.601** | 0.747** | 0.742** |
|  | SR | 0.782** | 0.779** | 10.000** | 1 | 0.722** | 0.559** | 0.110 | -0.183 | 0.716** | 0.686** | 0.654** | 0.591** |
|  | - | -0.059 | -0.012 | 9.021 | - | 0.065 | -0.113 | 0.275 | -0.368 | 0.046 | 0.085 | -0.093 | -0.151 |
| RL (cm) | FR | 0.836** | 0.783** | 0.683** | 0.657** | 1 | 0.987** | -0.408** | 0.187 | 0.942** | 0.971** | 0.964** | 0.930** |
|  | SR | 0.841** | 0.799** | 0.722** | 0.722** | 1 | 0.834** | 0.032 | -0.246 | 0.952** | 0.970** | 0.913** | 0.842** |
|  | - | 0.005 | 0.016 | 0.039 | 0.065 | - | -0.153 | 0.44 | -0.433 | 0.01 | -0.001 | -0.051 | -0.088 |
| RSA (cm<sup>2</sup>) | FR | 0.875** | 0.845** | 0.702** | 0.672** | 0.987** | 1 | -0.277* | 0.146 | 0.984** | 0.967** | 0.936** | 0.897** |
|  | SR | 0.754** | 0.706** | 0.559** | 0.559** | 0.834** | 1 | -0.012 | -0.333* | 0.813** | 0.786** | 0.753** | 0.689** |
|  | - | -0.121 | -0.139 | -0.143 | -0.113 | -0.153 | - | 0.265 | -0.479 | -0.171 | -0.181 | -0.183 | -0.208 |
| RAD (mm) | FR | -0.107 | 0.003 | -0.142 | -0.165 | -0.408** | -0.277* | 1 | -0.344* | -0.122 | -0.377** | -0.465** | -0.468** |
|  | SR | 0.098 | 0.137 | 0.110 | 0.110 | 0.032 | -0.012 | 1 | -0.550** | 0.314* | -0.022 | 0.047 | 0.059 |
|  | - | 0.205 | 0.134 | 0.252 | 0.275 | 0.44 | 0.265 | - | -0.206 | 0.436 | 0.355 | 0.512 | 0.527 |
| RLPV (cm/m<sup>3</sup>) | FR | 0.155 | 0.065 | 0.180 | 0.185 | 0.187 | 0.146 | -0.344* | 1 | 0.092 | 0.193 | 0.231 | 0.263 |
|  | SR | -0.323* | -0.348* | -0.183 | -0.183 | -0.246 | -0.333* | -0.550** | 1 | -0.400** | -0.195 | -0.164 | -0.152 |
|  | - | -0.478 | -0.413 | -0.363 | -0.368 | -0.433 | -0.479 | -0.206 | - | -0.492 | -0.388 | -0.395 | -0.415 |
| RV (cm<sup>3</sup>) | FR | 0.893** | 0.889** | 0.703** | 0.670** | 0.942** | 0.984** | -0.122 | 0.092 | 1 | 0.932** | 0.876** | 0.833** |
|  | SR | 0.846** | 0.809** | 0.716** | 0.716** | 0.952** | 0.813** | 0.314* | -0.400** | 1 | 0.901** | 0.871** | 0.801** |
|  | - | -0.047 | -0.08 | 0.013 | 0.046 | 0.01 | -0.171 | 0.436 | -0.492 | - | -0.031 | -0.005 | -0.032 |
| Tips | FR | 0.812** | 0.776** | 0.618** | 0.601** | 0.971** | 0.967** | -0.377** | 0.193 | 0.932** | 1 | 0.904** | 0.888** |
|  | SR | 0.811** | 0.764** | 0.686** | 0.686** | 0.970** | 0.786** | -0.022 | -0.195 | 0.901** | 1 | 0.866** | 0.821** |
|  |  | -0.001 | -0.012 | 0.068 | 0.085 | -0.001 | -0.181 | 0.355 | -0.388 | -0.031 | - | -0.038 | -0.067 |
| Forks | FR | 0.830** | 0.761** | 0.767** | 0.747** | 0.964** | 0.936** | -0.465** | 0.231 | 0.876** | 0.904** | 1 | 0.976** |
|  | SR | 0.756** | 0.737** | 0.654** | 0.654** | 0.913** | 0.753** | 0.047 | -0.164 | 0.871** | 0.866** | 1 | 0.965** |
|  |  | -0.074 | -0.024 | -0.113 | -0.093 | -0.051 | -0.183 | 0.512 | -0.395 | -0.005 | -0.038 | - | -0.011 |
| Crossings | FR | 0.792** | 0.718** | 0.766** | 0.742** | 0.930** | 0.897** | -0.468** | 0.263 | 0.833** | 0.888** | 0.976** | 1 |
|  | SR | 0.689** | 0.684** | 0.591** | 0.591** | 0.842** | 0.689** | 0.059 | -0.152 | 0.801** | 0.821** | 0.965** | 1 |
|  |  | -0.103 | -0.034 | -0.175 | -0.151 | -0.088 | -0.208 | 0.527 | -0.415 | -0.032 | -0.067 | -0.011 | - |

**Fig. S1.**
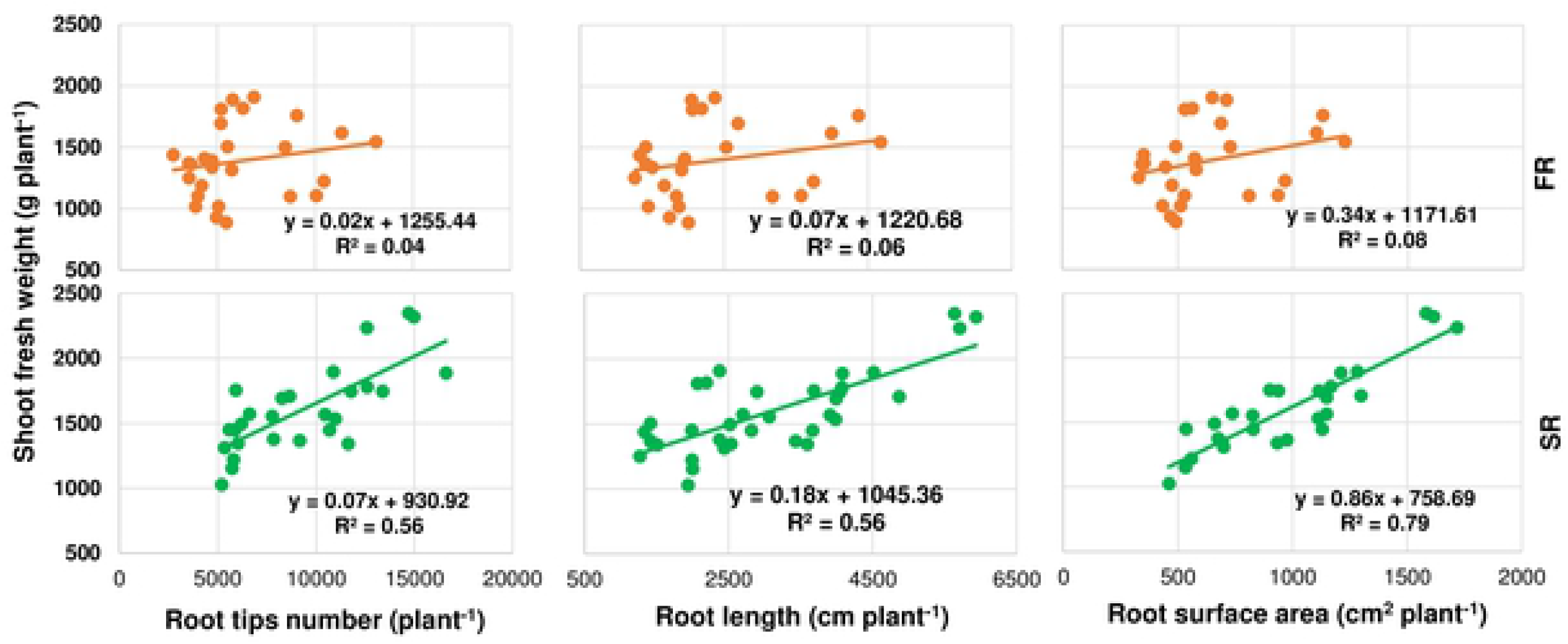
Relationships between shoot biomass and root length, root tip number, and root surface area in the first (FR) and second ratoon crop (SR). Plant^−1^: the amount per each cluster / millable canes in the cluster.

